# High-resolution TADs reveal DNA sequences underlying genome organization in flies

**DOI:** 10.1101/115063

**Authors:** Fidel Ramírez, Vivek Bhardwaj, José Villaveces, Laura Arrigoni, Björn A. Grüning, Kin Chung Lam, Bianca Habermann, Asifa Akhtar, Thomas Manke

**Affiliations:** Max Planck Institute of Immunobiology and Epigenetics. Bioinformatics Unit. Stübeweg 51, 79108 Freiburg Germany; Max Planck Institute of Biochemistry. Computational Biology. Am Klopferspitz 18, 82152 Martinsried Germany; Max Planck Institute of Immunobiology and Epigenetics. Sequencing Facility. Stübeweg 51, 79108 Freiburg Germany; University of Freiburg, Department of Computer Science, Georges-Köhler-Allee 106, 79110 Freiburg, Germany; Faculty of Biology, University of Freiburg, SchänzIestraße 1, 79104 Freiburg, Germany

## Abstract

Eukaryotic chromatin is partitioned into domains called TADs that are broadly conserved between species and virtually identical among cell types within the same species. Previous studies in mammals have shown that the DNA binding protein CTCF and cohesin contribute to a fraction of TAD boundaries. Apart from this, the molecular mechanisms governing this partitioning remain poorly understood. Using our new software, HiCExplorer, we annotated high-resolution (570 bp) TAD boundaries in flies and identified eight DNA motifs enriched at boundaries. Known insulator proteins bind five of these motifs while the remaining three motifs are novel. We find that boundaries are either at core promoters of active genes or at non-promoter regions of inactive chromatin and that these two groups are characterized by different sets of DNA motifs. Most boundaries are present at divergent promoters of constitutively expressed genes and the gene expression tends to be coordinated within TADs. In contrast to mammals, the CTCF motif is only present on 2% of boundaries in flies. We demonstrate that boundaries can be accurately predicted using only the motif sequences, along with open chromatin, suggesting that DNA sequence encodes the 3D genome architecture in flies. Finally, we present an interactive online database to access and explore the spatial organization of fly, mouse and human genomes, available at http://chorogeome.ie-freiburg.mpg.de.

## Introduction

The partitioning of chromosomes into topologically associating domains (TADs) is an emerging concept that is reshaping our understanding of gene regulation in the context of physical organization of the genome (Dixon et al. 2016; Lieberman-Aiden et al. 2009; Rao et al. 2014; Sexton et al. 2012; Hou et al. 2012). However, the mechanisms by which chromatin acquires its 3-dimensional conformation are not fully understood. TADs were first described in mouse ESC cell lines (Nora et al. 2012) as regions of preferential DNA contact. Since then, TADs have been discovered in other mouse and human cell lines (Dixon et al. 2012) as well as in other mammals (dog and macaque (Vietri Rudan et al. 2015), in flies (Sexton et al. 2012; Ramírez et al. 2015;Hou et al. 2012), worms (Crane et al. 2015) and yeast (Mizuguchi et al. 2014). TAD-like structures have also been reported in plants (Arabidopsis thaliana) (Wang et al. 2015) and bacteria (*Caulobacter crescentus*) (Le et al. 2013). Interestingly, TADs seem to be virtually identical between cell lines of the same organism (Dixon et al. 2012; Ramírez et al. 2015; Rao et al. 2014) and they are conserved between species (Vietri Rudan et al. 2015). Recent publications show that TADs correspond to cytogenetic bands (Ulianov et al. 2015; Eagen et al. 2015). The functional importance of domains and their proper separation was highlighted by (Lupiáñez et al. 2015) who demonstrated that disruption of TAD boundaries cause changes in gene-enhancer interactions that lead to developmental abnormalities in mouse embryos. This is consistent with the idea that TADs compartmentalize regulation by inhibiting enhancer-promoter interactions with neighboring TADs (Smallwood & Ren 2013).

To understand TAD formation, researchers had focused on the proteins found at TAD boundaries (Dixon et al. 2012; Sexton et al. 2012; Hou et al. 2012). In mammalian cells, the the CCCTC-binding factor (CTCF) protein has been shown to be enriched at chromatin loops, which also demarcate a subset of TAD boundaries (referred to as “loop domains”) (Rao et al. 2014). A proposed mechanism, based on the extrusion of DNA by cohesin, suggests that the DNA binding motif of CTCF and its orientation determine the start and end of the loop (Sanborn et al. 2015; Nichols & Corces 2015). In line with this hypothesis, deletions of the CTCF DNA-motif effectively removed or altered the loop (Sanborn et al. 2015). Interestingly, mutations of the CTCF binding motif at TAD boundaries are particularly abundant in cancer, causing dysregulation of oncogenes (Hnisz et al. 2016). However, CTCF-cohesin loops only explain a fraction (less than 39%) of human TADs boundaries (Rao et al. 2014), while plants and bacteria lack CTCF homologs but also show TAD-like compartments. Thus, it is possible that additional proteins are involved in the formation of TADs.

In contrast to mammals, the genetic manipulation tools available in flies have allowed the characterization of several proteins that, like CTCF, are capable of inhibiting enhancer-promoter interactions. Throughout the manuscript, we will refer to these proteins as “insulator proteins” and their binding motifs as “Insulators” or “Insulator motifs”. In flies, apart from CTCF, the following DNA-binding insulator proteins have been associated to boundaries (Sexton et al. 2012; Van Bortle et al. 2014): Boundary Element Associated Factor-32 (Beaf-32), Suppressor of Hairy-wing (Su(hw)) and GAGA factor (GAF). Also, Zest white 5 (Zw5) has been proposed to bind boundaries (Zolotarev et al. 2016a). These insulator proteins recruit co-factors critical for their function such as Centrosomal Protein-190 (CP190) and Mod(mdg4) (Ong & Corces 2014). Recently, novel insulators have been described as binding partners of CP190: the zinc finger protein interacting with CP190 (ZIPIC), Pita (Maksimenko et al. 2015) which appear to have human homologs and localizes to TAD boundaries (Zolotarev et al. 2016a), and the Insulator binding factors 1 and 2 (Ibf1 and Ibf2) (Cuartero et al. 2014). All the previously characterized boundary associated proteins bind to the DNA at specific motifs, suggesting that the 3D conformation of chromatin can be encoded by these motifs.

In this study, we sought to identify the DNA encoding behind TADs boundaries in flies. First, we developed new software (HiCExplorer) to obtain boundary positions at 0.5 kilobase resolution based on published Hi-C sequencing data from *Drosophila melanogaster* Kc167 cell line (Li et al. 2015; Cubeñas-Potts et al. 2016). Using these high-resolution TAD boundaries we identify eight significantly enriched DNA-motifs. Reassuringly, five of these motifs are known to be bound by the insulator proteins: Beaf-32, CTCF, the heterodimer Ibf1 and Ibf2, Su(Hw) and ZIPIC. The three remaining DNA-motifs have not been associated to boundaries before. Interestingly, one of the motifs is bound by the motif-1 binding protein (M1BP) (Li & Gilmour 2013), a protein associated to constitutively expressed genes whose role as insulator remains unexplored.

Using machine learning methods based on the acquired DNA motif information, we could accurately distinguish boundaries from non-boundaries and identify new TAD boundaries that were missed when using only Hi-C data. This suggests that the chromosomal folding in flies can be explained predominantly by the DNA sequence alone.

We implemented the methods for Hi-C data processing, TAD calling and visualization into an easy to use tool called HiCExplorer (hicexplorer.readthedocs.io). To facilitate exploration of available Hi-C data, we also provide an interactive online database containing processed high-resolution Hi-C data sets from fly, mouse and human genome, available at http://chorogeome.ie-freiburg.mpg.de.

## Results

### High resolution TAD boundaries in flies

We obtained Hi-C data for Kc167 cells from (Li et al. 2015; Cubeñas-Potts et al. 2016) and processed them to obtain corrected Hi-C contact matrices at restriction fragment resolution (see methods). These datasets contain the most detailed contact maps in flies, compared to other datasets (see methods), due to high sequencing depth (over 246 million valid read pairs) and the use of DpnII, a restriction enzyme with short restriction fragment size (mean ~570 bp). We found 2,852 TADs having a median size of 26 kb (Fig. 1A, S1A). We corroborated the precision of our boundaries by comparing their overlap with CP190 peaks (*p*-value = 1.8E-20, Fig. 1B) and the separation of histone marks (Fig. 1C, S1B). Similar to (Sexton et al. 2012), we classified TADs using modENCODE histone marks as active (enriched for either H3K36me3, H3K4me3 and H4K16ac), polycomb group silenced (PcG) (enriched for H3K27me3), HP1 (enriched for H3K9me3) and inactive (not enriched for any of the marks, see methods and Fig. S1C). A significant fraction of the genome (43%) is covered by large inactive TADs having a mean length of 63 kb (Fig. 1D). In contrast, active chromatin TADs have an average length of 23 kb and occupy 29% of the genome. PcG chromatin occupies 25% of the genome with TADs that are on average 61 kb. The largest TADs are found for HP1 repressed chromatin, which occupies each 3% of the genome and have a length of 74 kb. We also find that active TADs tend to be assembled one after the other due to their higher number (Fig. 1E). Interestingly, the TAD separation score varied significantly (*p*-value < 7.8E-5, Wilcoxon rank-sum test) between the TAD types (Fig. 1F). The stronger boundaries (low TAD-separation score) are found between active and inactive or PcG TADs. while the weakest boundaries are found between PcG TADs. Similarly, we find that the TAD-separation score between larger TADs (mostly inactive) is significantly larger than the TAD separation score for smaller TADs (mostly active) (*p*-value = 9.9E-7, Wilcoxon rank-sum test).

**Figure 1.**
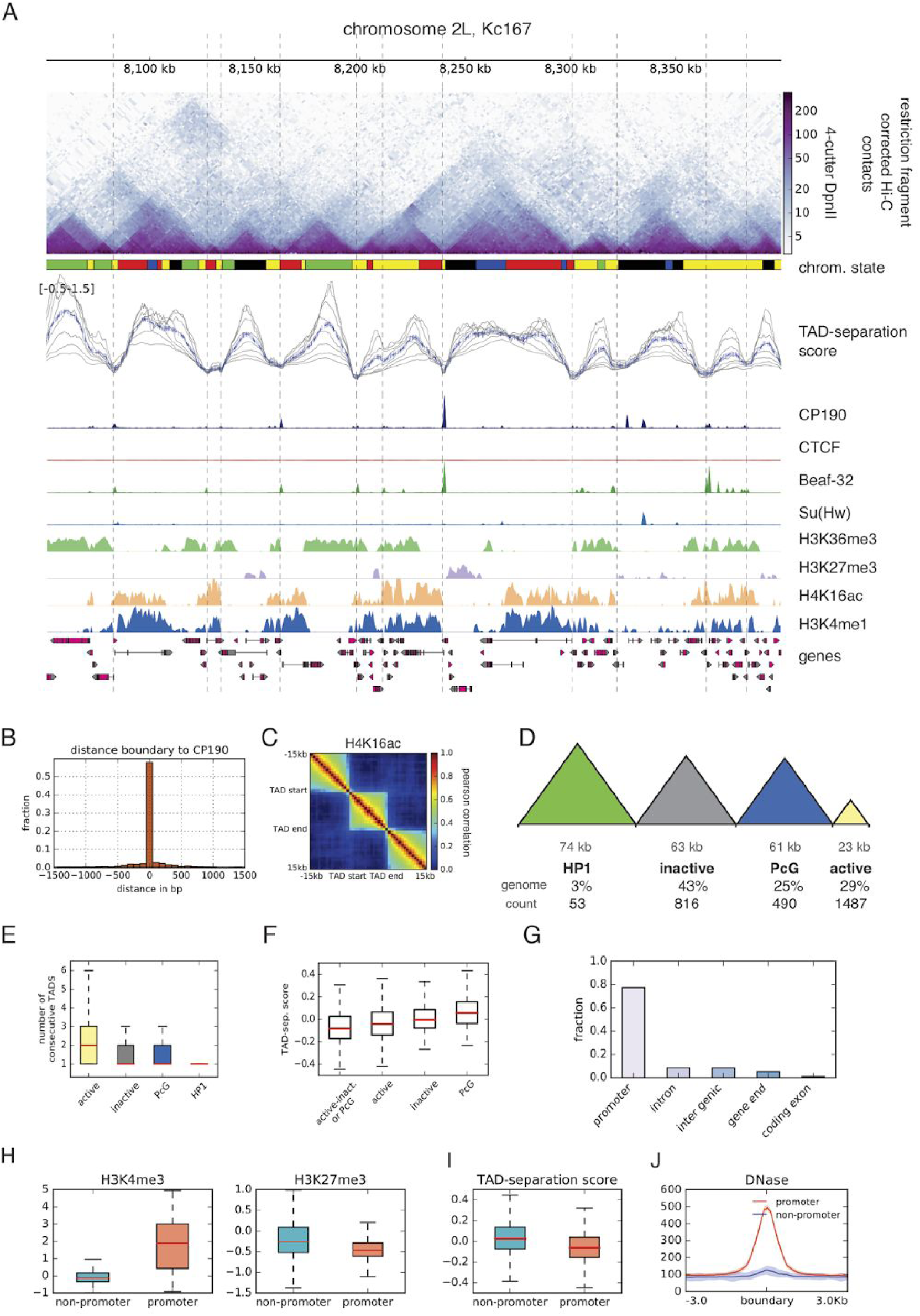
High-resolution TAD boundaries in flies. **A.** Example region of 350 kb showing Hi-C TADs from Kc167 cells. The top panel shows a heatmap of corrected counts from the Hi-C contact matrix obtained from (Li et al. 2015; Cubeñas-Potts et al. 2016). The size of the bins is variable (mean 570 bp) and depends on the genomic location of the DpnII restriction sites. The chromatin state track contains the five classifications from (Filion et al. 2010): Active chromatin, red and yellow; inactive chromatin, black; PcG, blue; HP1, green. The TAD separation score track (see methods) depicts a normalized measure of the contacts between two flanking regions. The gray lines show scores for flanking regions of different size ranging from 10 kb to 40 kb and the blue line shows the mean score. The TAD-separation score was used to identify local minima indicative of TAD boundaries. The estimated boundaries are shown as vertical lines. The following four tracks show normalized ChIP-seq coverage for the insulators CP190, Beaf-32, and Su(Hw) on Kc167 from (Wood et al. 2011) and CTCF from (Li et al. 2015) that are known to be associated to boundaries (Sexton et al. 2012; Van Bortle et al. 2014). The following tracks contain ChIP-chip data for histone modifications from modEncode (Celniker et al. 2009). This particular region was selected because many different TADs could be seen; other regions can be browsed at http://chorogenome.ie-freiburg.mpg.de:5001. **B.** Histogram of the distance of a boundary to the nearest CP190 (common insulator protein co-factor) peak. **C.** Correlation of histone marks within and between TADs. Each pixel in the matrix represent the pearson correlation of the histone mark in all TADs at different distances (see methods). **D.** TAD classification based on histone marks. The numbers below each TAD type represent respectively: mean length, percentage of genome occupied by the TAD and number of TADs of that type. **E.** Boxplot of consecutive TAD of each type. **F.** TAD-separation score between: active and inactive or PcG, active-active, inactive-inactive and PcG-PcG. The differences between the groups are all significant (*p*-value < 7.8E-5, Wilcoxon rank-sum test). **G.** Classification of TAD boundaries. TAD boundaries are classified at promoter if they are within 1000 bp of the annotated TSS. **H.** Histone marks at non-promoter and promoter boundaries. Further marks can be seen in Fig. S1I. **I**. TAD-separation score for non-promoter and promoter boundaries (*p*-value=8.52E-35, Wilcoxon rank-sum test). **J.** DNase accessibility at non-promoters and promoter boundaries.

While most of our boundaries overlap with those from previous studies (Sexton et al. 2012; Cubeñas-Potts et al. 2016), our method allowed us to identify a larger set of boundaries (Fig. S1D-H) which mostly divide active TADs from previous studies. We observed that the majority of the boundaries (77%) are located at gene promoters (henceforth referred to as promoter-boundaries. Fig. 1G). Promoter-boundaries are different from non-promoter boundaries (27%), since they associate significantly with active chromatin (Fig. 1H), have lower TAD separation score (representing strong boundaries Fig. 1I), and show higher DNAse sensitivity (Fig. 1J).

### Boundaries are marked by specific gene orientation and transcription

Next, we correlated our promoter boundaries with gene orientation and transcription. Most promoter-boundaries (70%, *p*-value = 2.5E-88 fisher's exact test), are associated with divergently oriented genes while genes in convergent or tandem orientation tend to be inside the TADs. To correlate TADs with gene transcription, we analysed the RNA-Seq data of 14 stages of Drosophila development along with the expression in the Kc167 cell line obtained from modENCODE (Fig. S2A, see methods). We found that, in general, 95.6% of genes associated with TAD boundaries are expressed in Kc167 cells compared to only 75.3% of genes which don’t have a boundary (*p*-value = 2.19E-80, Fisher’s test). Higher correlation was observed between gene expression inside TADs than between neighboring TADs (Fig. 2A, S2C). When we investigated the expression of genes at TAD boundaries we found that these genes show significantly higher expression than the expressed genes which don’t have a TAD boundary at their promoters (Fig 2B, *p*-value = 6.08E–21, t-test). Boundary associated genes also show a more stable expression expression across development than other genes (Fig 2C, S2B), suggesting that these genes are ubiquitously transcribed. Furthermore, we find that for a pair of genes lying next to each other, the variability in their gene expression tend to be correlated during development if these genes are within same TAD, while this correlation is lost if there is a TAD boundary in between (Fig 2D, see methods). This is true for gene-pairs in any orientation (convergent, divergent or tandem, Fig S2D).

**Figure 2.**
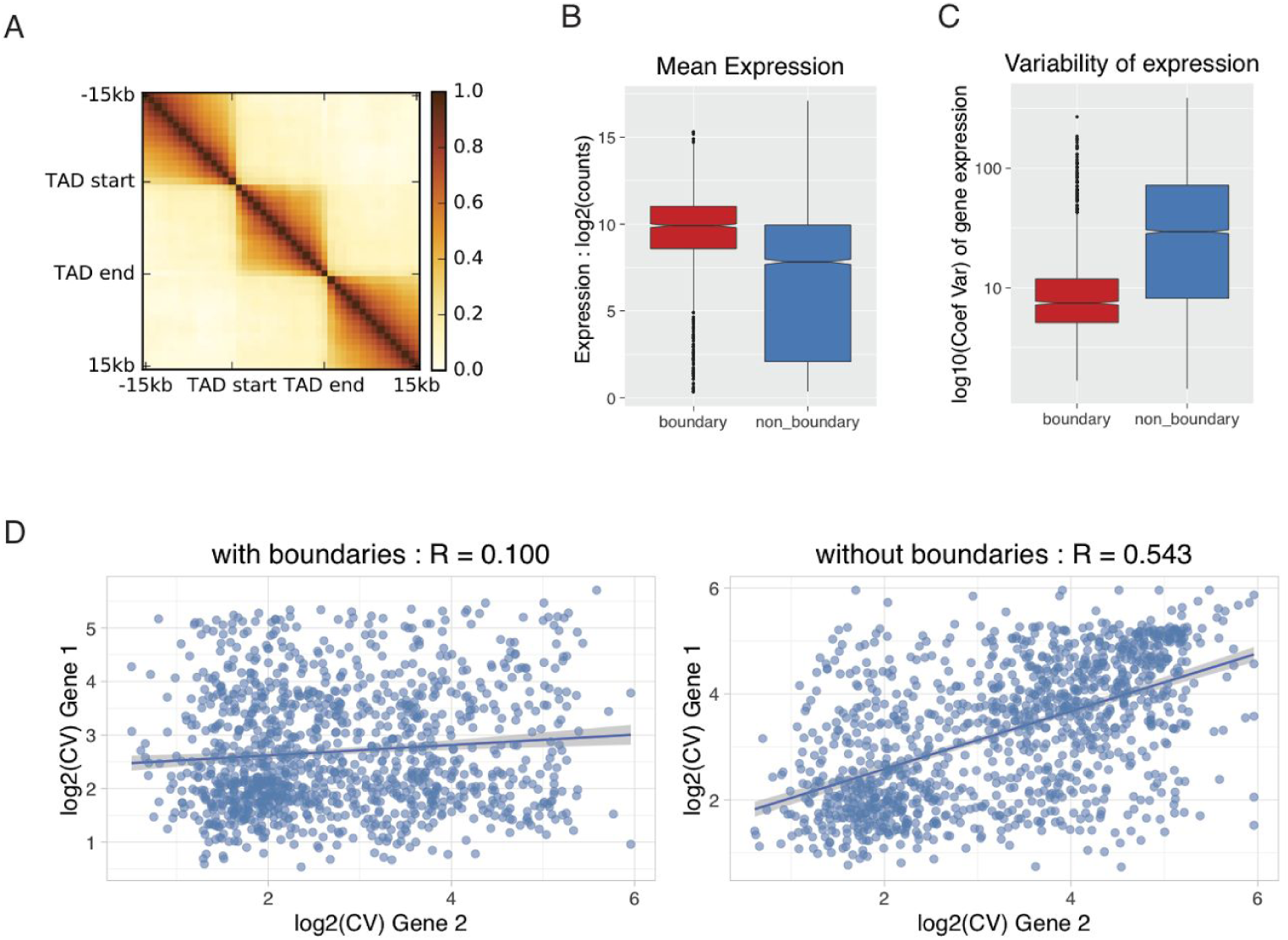
TAD boundaries are marked by specific gene orientation and transcription A. Correlation of mean expression across developmental stages inside TADs vs outside. Region inside TADs was scaled to 15 kb. Each pixel in the matrix contains the pearson correlation at different distances. **B-C.** Mean expression (in Kc167 cells) and variability of expression (during development) for genes whose promoters are at a TAD boundary vs. genes whose promoters are not at boundaries. **D.** Coefficient of variation (across developmental stages) between pairs of adjacent genes either separated by a TAD boundary (left) or not separated by a boundary (right).

Taken together, these results suggest that specific gene orientation and level of transcription could be associated with TAD formation.

### A comprehensive list of boundary associated DNA motifs

We followed the strategy outlined in (Fig 3A) to create a comprehensive list of motifs frequently found at boundaries. First, we performed *de-novo* motif calling using MEME-chip (Ma et al. 2014) (see methods) on our promoter-boundaries and non-promoter boundaries. To filter out motifs that are frequently found at promoters or open chromatin but are not specific to boundaries, we did an enrichment analysis using two different methods: Ame (McLeay & Bailey 2010) and TRAP (Thomas-Chollier et al. 2011). As a second approach, we tested boundaries for the motifs of known insulator proteins and core-promoter motifs from (Ohler et al. 2002). In contrast to *de-novo* motif detection, searching for known motifs allows additional sensitivity to detect low frequency motifs. After filtering for only consistent results, we could identify 5 motifs enriched at promoter-boundaries and 3 motifs enriched at non-promoter boundaries (Fig 3B).

**Figure 3.**
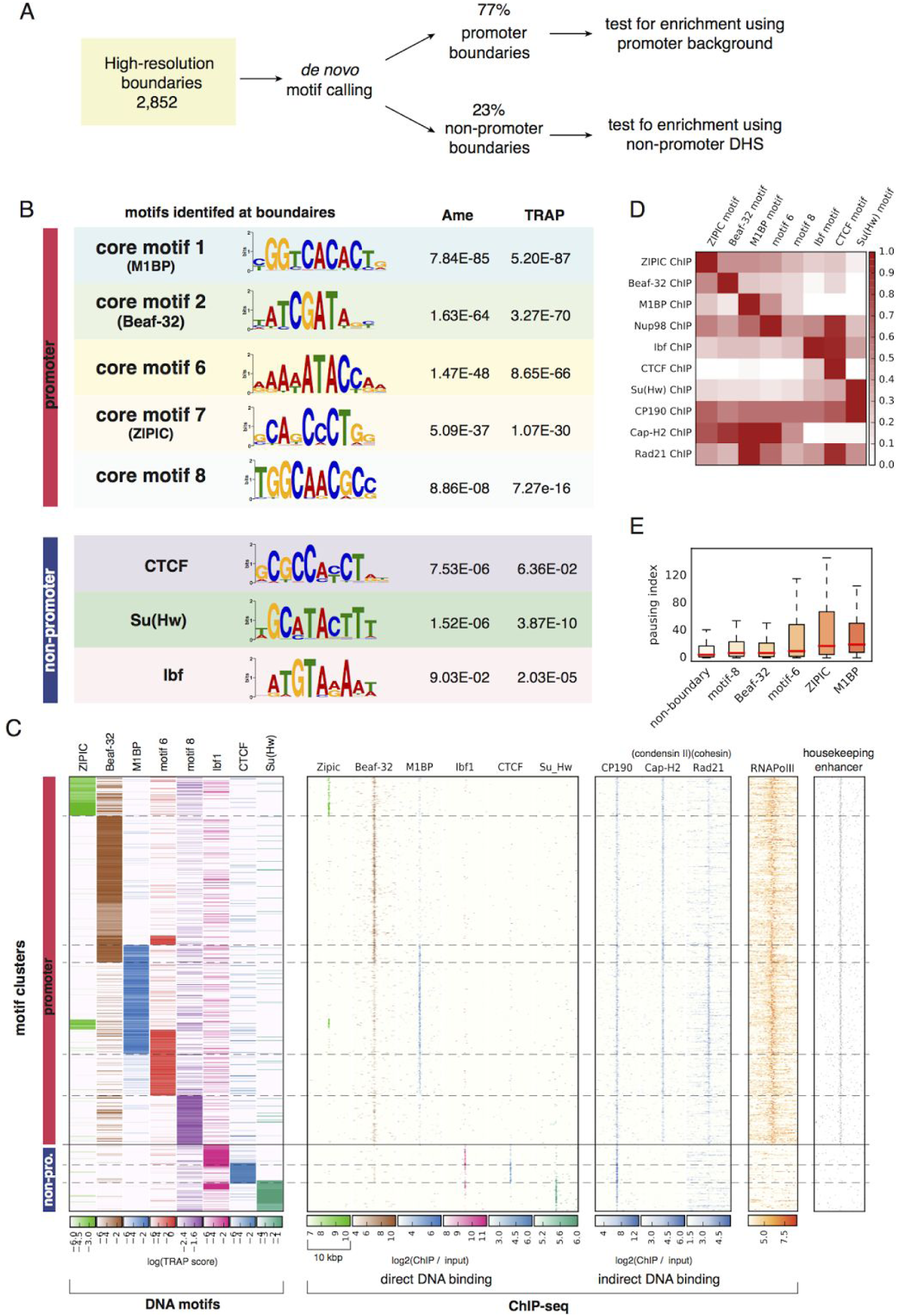
Eight motifs are enriched at boundaries: A. Overview of the strategy used to identify *de-novo* motifs. **B.** Motifs enriched at promoter and non-promoter boundaries (along with corrected *p*-values). Two methods were used to estimate enrichment (see methods): Ame (McLeay & Bailey 2010) and TRAP (Thomas-Chollier et al. 2011). **C.** Clustering of boundaries by motif binding affinity (see methods). Each row represents one boundary. Left panel: clustering of motif binding affinity using the TRAP score (Thomas-Chollier et al. 2011). Higher scores indicate stronger predicted binding. Dashed lines delineate the clusters. Following panels: Using the motif clustering results we show the heatmaps corresponding to ChIP-seq enrichments for insulators protein binding the DNA (second panel), other proteins that bind indirectly (third panel) and RNA Pol-II. Last panel shows housekeeping enhancers from (Zabidi et al. 2015). For boundaries at promoters, heatmaps are centered at the gene promoter, for non-promoter boundaries, heatmaps are centered at the nearest CP190 peak within 2000 bp. ChIP-seq signal was computed in 50 bp bins for 5000 bp from the center. The scale of each heatmap was adjusted based on the distribution of the TRAP or ChIP-seq values in the respective cluster (Fig. S3 A and B). **D.** Relationship between motif presence and ChIP-seq peak fold change at boundaries. Each cell in the matrix contains the mean fold change of all respective ChIP-seq peaks having the motif. For each row, the maximum fold change was scaled to 1. **E.** Pausing index at different boundary-promoters containing one of the boundary motifs. Non-boundary promoters are plotted as control.

The promoter boundary motifs we identified belong to the list of core-promoter motifs 1, 2, 6, 7 and 8 from (Ohler et al. 2002). Motif-1 is recognized by the recently described 'motif-1 binding protein' (M1BP), a protein found at the promoters of transiently paused Pol-II of constitutively expressed genes (Li & Gilmour 2013). Motif-2 (also called DRE motif) is recognized by the insulator protein Beaf-32 and DREF (Gurudatta et al. 2013). Motif-7 (also called DMv3) is recognized by the insulator protein ZIPIC. The binding proteins for motif-6 (also known as DMv5) and motif-8 (also known as DMv2) are, to the best of our knowledge, not known. *De-novo* motif calling also identified other core-promoter motifs but they were not found enriched at boundaries. The three motifs that we find enriched on non-promoter boundaries, correspond to the binding sites of Su(Hw) and CTCF and Ibf (Fig 3B). We could not find an enrichment for motifs of other insulator proteins like GAF, Pita and Zw5. For clarity, we will refer to the boundary motifs by the name of the insulator protein that binds to them, except for motif-6 and 8.

As an independent validation, we find that our three novel boundary motifs (M1BP motif, motif-6 and motif-8) are also enriched at the binding site of CP190, Rad21 (part of cohesin complex) and Cap-H2 (condensin II complex) (Table S1). Additionally, we repeated our analysis using the TAD boundaries from (Sexton et al. 2012) and (Li et al. 2015; Cubeñas-Potts et al. 2016) and found similar enrichments (Table S2).

To better understand the distribution of the motifs on boundaries we performed hierarchical clustering of the binding affinity (TRAP score) for the eight motifs enriched at boundaries (Fig. 3C left panel). We then plotted the ChIP-seq signal of the DNA binding proteins (Fig. 3C second panel), along with CP190, Cap-H2, Rad21 (Fig. 3C third panel) and RNA Pol-II, over the clusters. The results show that the boundary motifs are usually associated with their corresponding proteins, except for motifs 6 and 8 for which the binding proteins are not known (Fig 3C second panel). Examples of the boundaries with their motifs and corresponding proteins can be seen in Fig. S3C-G.

We discovered that in addition to the Beaf-32 motif, a novel M1BP motif is most frequently found at boundaries. In combination those two motifs are enriched at 55% of promoter boundaries, and are associated with their corresponding proteins. Promoter boundaries also tend to be associated with condensin II (Cap-H2), cohesin (Rad21), RNA polymerase II and housekeeping enhancers. On the other hand Ibf1, CTCF and Su(Hw) are the most common proteins associated with non-promoter boundaries, and tend to be associated with enhanced binding of CP190 co-factor (Fig. 3C, 3D). Similar to this result, we observe that the binding sites of promoter and non-promoter boundary proteins are correlated (Fig. S3H).

Since M1BP binds paused RNA Pol-II promoters (Li & Gilmour 2013), we checked whether other promoter-boundary motifs are also associated with paused RNA Pol-II. Indeed, we find that besides M1BP, promoters containing the ZIPIC motif or motif 6 also associate with paused Pol-II (Fig. 3E).

We then searched for association of all the transcription factors from modENCODE consortium (Celniker et al. 2009) as well as from the comprehensive collection of ChIP-seq data from (Li et al. 2015; Cubeñas-Potts et al. 2016) with our boundary motifs. Similar to Fig. 3C, all motifs are enriched for binding sites of their corresponding proteins. Interestingly, our screen for proteins associated to boundary motifs showed that Nup98, a component of the nuclear pore complex, is associated with motif-6 and Pita (Fig. 3D and S3I). Further validation using *de-novo* motif calling on Nup98 peaks and identified motif 6 and Pita motif as the most enriched motifs (q = 3.4E-4 and 3.3E-09, respectively).

We observed that ChIP-seq peaks of DNA binding proteins are often found in regions not containing their motif (Fig. 3D). For example, ZIPIC peaks can be seen together with motif 6 or CTCF although ZIPIC motif does not overlap with any of them (Fig. 4A). Similar observations can be made for CTCF ChIP-seq experiments (Fig. S3J, see discussion), suggesting that motif sequences should be considered along with ChIP-Seq binding sites as more reliable functional predictors of an insulator protein.

**Figure 4.**
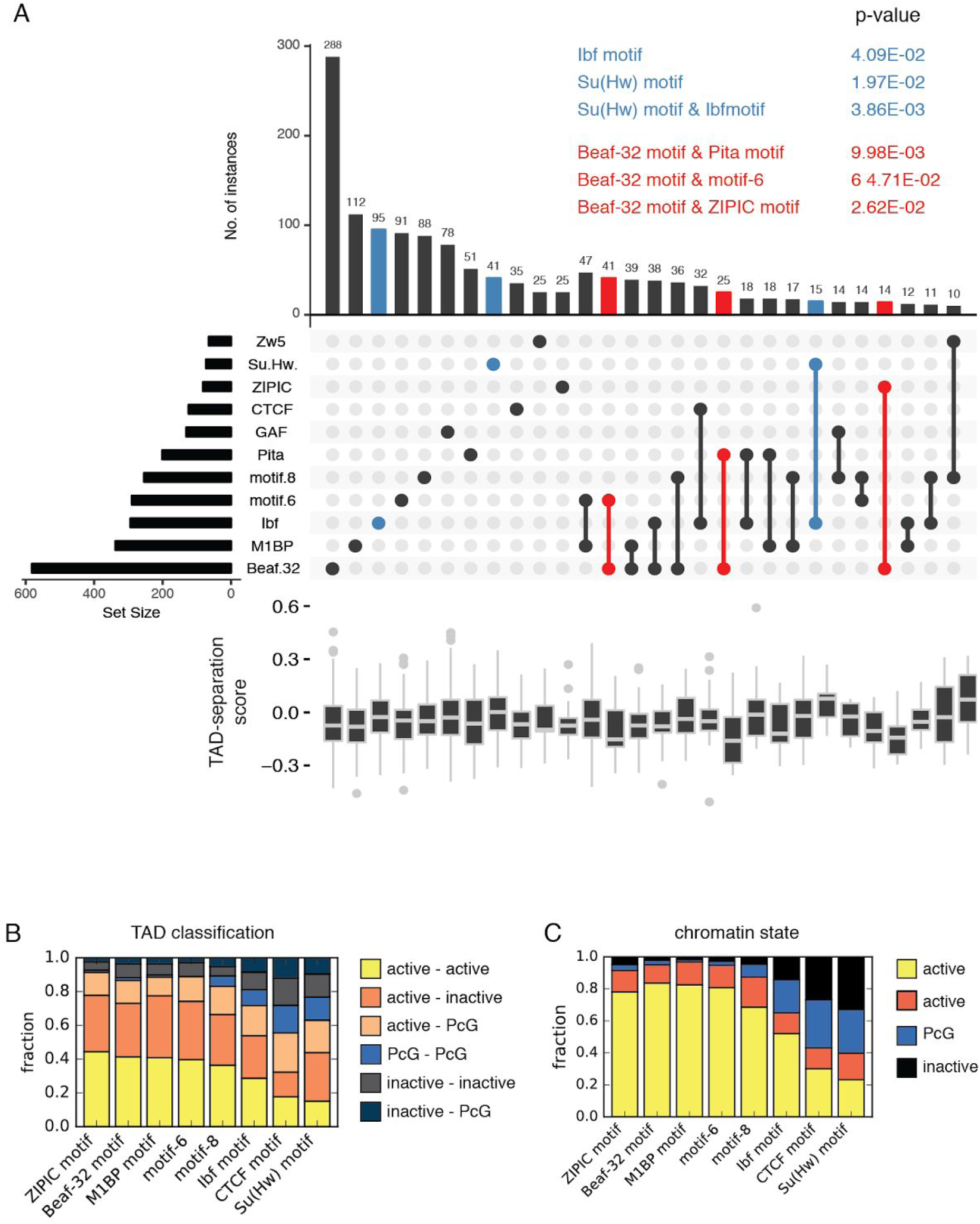
Promoter and non promoter motifs show marked differences. **A.** TAD-separation score at boundaries grouped by the motif presence. For this analysis we considered a motif to be present if the motif overlaps with a ChIP-seq peak (see methods). The bars show the overlap between the indicated motifs below. The boxplots show the distribution of the respective TAD-separation score. The sets highlighted in blue have a TAD-separation score distribution significantly larger than the overall TAD-separation score. The *p-*values (Wilcoxon rank-sum test) are show above the figure. Similarly, the sets highlighted in red have a distribution significantly smaller. Only motif combinations having more than 10 instances are shown. Motif combinations with three or more motifs were rare. The intersections were plotted using UpSetR (Lex et al. 2014). An overview of the motif overlaps can be seen in Fig. S4A. **B.** Frequency of flanking TAD types (as classified in Fig. 1D) per boundary motif. **C.** Frequency of the chromatin state from (Filion et al. 2010).

### Motif combinations reflect boundary strength and chromatin types

Next we looked at motif combinations at boundary, in relation to boundary strength. Using ChIP-Seq analysis, it has been reported that boundary strength increases with the number of proteins bound at the boundary (Van Bortle et al. 2014). When we looked for motif combinations at boundaries that overlap with their corresponding protein, we we didn’t observe any significant differences in boundary strength between boundaries containing one, two or three motifs (we only found a handful of boundaries having more than three motifs). Instead, we found that specific motifs or their combinations are associated with boundary strength (Fig. 4A). Boundaries containing the motif for Ibf, Su(Hw) or the combination of the two motifs are weaker than average while the combination of the Beaf-32 motif with either Pita, ZIPIC or motif-6 result in the strongest boundaries.

We also looked at the association of motifs with active, inactive, PcG and Hp1 TADs from Fig. 1D. We observe that the promoter-boundary motifs (Fig 4B) are mostly found between active TADs or between active and inactive (including PcG) TADs. Conversely, the non-promoter boundary motifs are rarely found between active TADs, and mostly separate active from inactive TADs or are between inactive TADs (Fig. 4B). We find the same trend when analysing the ChIP-chip intensity of the active histone mark H3K36me3 and the repressive histone mark H3K27me3 surrounding the boundaries (Fig. S4A). Most promoter-boundary motifs separate active-inactive marks, while most non-promoter boundary motifs lie within inactive marks.

Additionally, we analyzed the chromatin state (Filion et al. 2010) that overlaps with the boundary motifs and found that promoter-boundary motifs lie mostly within active chromatin while non-promoter boundary motifs lie within both active and inactive chromatin (Fig. 3C).

### Boundaries can be predicted using motifs

To better characterize boundaries at promoters we used three standard classification methods and ranked features by their relevance to distinguish boundaries from other promoters (see methods). The features included the TRAP score of all motifs studied along with other known insulator motifs. We also used DNase hypersensitive sites (DHS) as a feature to identify open promoters, considering that protein binding to a motif requires an accessible promoter. The ranking of feature importance showed that indeed open promoters are required for boundaries. For the motifs, the feature importance ranking follows the abundance of the motifs at boundaries: Beaf-32 and M1BP appear to be most important, followed by motif-6, ZIPIC and motif-8. Some features, like GAF and Pita were found to be negatively associated with boundaries at promoters (Fig. S5A).

Although regularized models used here are less prone to overfitting compared to other machine learning methods, we further protect against overfitting by 10-fold cross validation during training. We then tested model accuracy on an independent test dataset. The predicted data from the classifiers showed good results, with a sensitivity and specificity over 71% (Fig. 5B). For the small fraction of non-promoter boundary motifs, we used open chromatin regions distant from promoters to train our classifiers using the non-promoter boundary motifs as features. We achieved prediction accuracy similar to the promoter-boundaries (Fig. 5C,D). Interestingly, we find GAF to be negatively associated with both promoter and non-promoter boundaries (Fig S5C).

**Figure 5.**
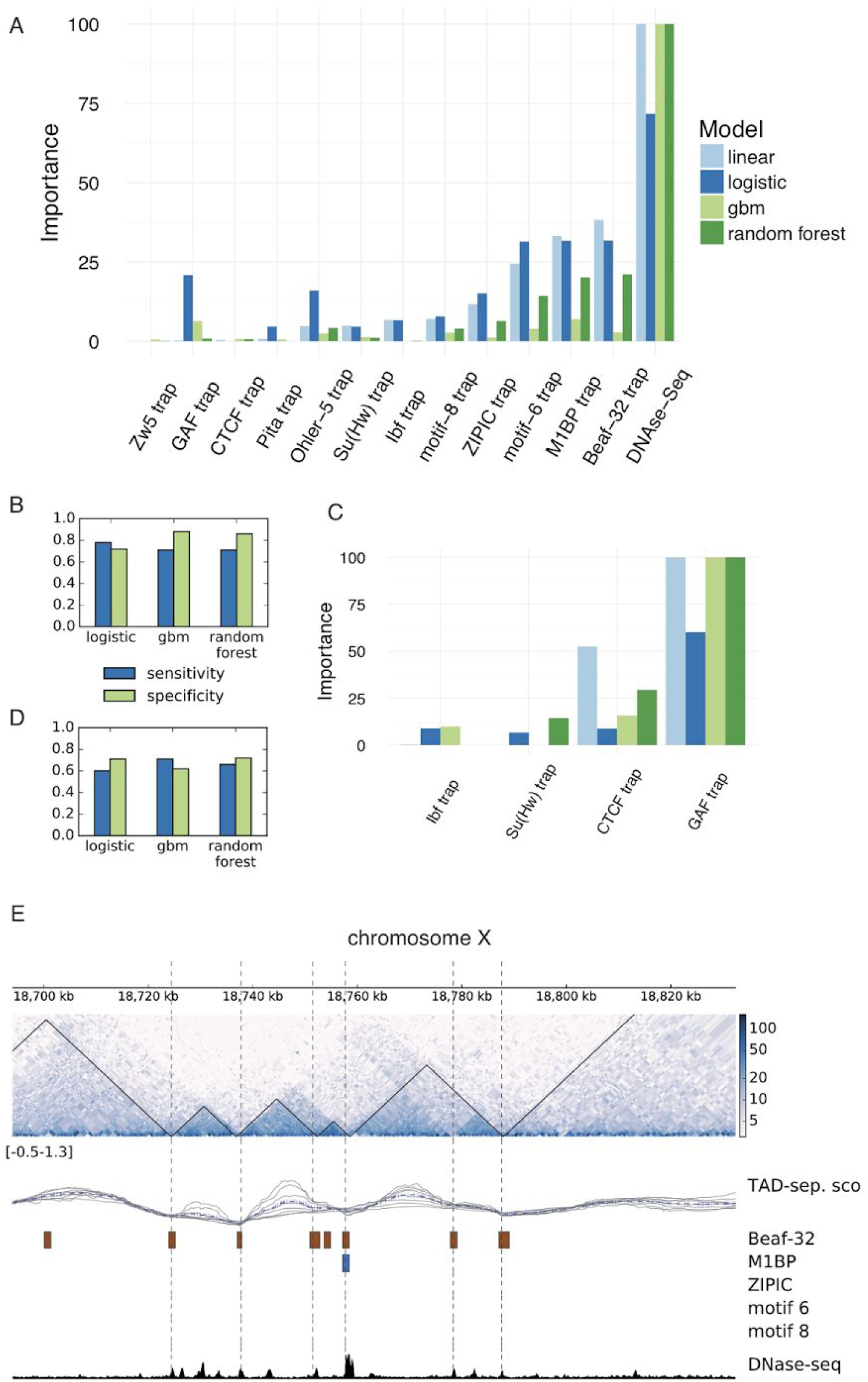
Machine learning prediction of boundaries. **A.** Feature importance for promoter boundaries computed using four different methods: linear model, logistic regression, gradient boost model (gbm) and random forest. Importance scores of each method were scaled from 0 to 100. Except for DNAse-seq, each feature represents the binding affinity (TRAP score) of the respective motif. **B.** Sensitivity and specificity for promoter boundaries measured for logistic regression, gradient boost model and random forest. The output of the linear model can be seen in Supplementary Fig. S3B. **C.** Feature rankings, as in (A), for non-promoter boundaries. **D.** Sensitivity and specificity for non-promoter boundaries. **E.** Examples of high-resolution boundaries and predicted boundaries. The high-resolution boundaries (based on the TAD-separation score) are depicted as black triangles on top of the Hi-C heatmap. The predicted boundaries are shown as dotted vertical lines. The tracks below the the Hi-C contact map contain the instances of the motifs that overlap with promoters. To aid the visualization of the short motifs, their genomic location was extended by 500 bp in each direction. The last track depicts regions of open chromatin based on DNase-seq from modEncode (Celniker et al. 2009).

We could evidence that the machine learning predictions can complement the Hi-C derived boundaries at the promoters, as some boundaries predicted by our model are missed by our boundary detection method based on the TAD-separation score. The predictions also detect putative boundaries that are very close to each other and can not be resolved using Hi-C data (Fig. 5E).

### Resources to identify and explore TADs and associated genomic features

During our research we developed processing and analysis tools for chromosome conformation. Our tool-suite, called HiCExplorer, simplifies the Hi-C data pre-processing, contact matrix transformation, and TAD calling into a few easy steps. HiCExplorer is open source and and available at https://github.com/maxplanck-ie/HiCExplorer. Importantly, HiCExplorer can be used with other pipelines and processing tools as we have built-in import/export functions covering commonly used Hi-C data formats. To facilitate analysis, we have integrated HiCExplorer into the Galaxy platform (Goecks et al. 2010). With HiCExplorer we made available our efforts to create meaningful and accurate visualizations of Hi-C data with other data sources, whose examples can be seen throughout this manuscript. Further information can be found at the associated documentation (http://hicexplorer.readthedocs.io), which includes a full analysis workflow and detailed description of the tools.

Since most users will not routinely perform an expensive and technically challenging Hi-C experiment, it will be highly beneficial to be able to visualize their genes or regions of interest along with associated TAD boundaries, to understand gene regulation. We provide a resource called the Chorogenome (http://chorogenome.ie-freiburg.mpg.de/), which includes Drosophila, Mouse and Human Hi-C datasets, already processed by HiCExplorer, along with associated gene annotations, histone marks and other TAD/boundary annotations. This can be used to quickly visualize any gene or region in context of TADs. The underlying program called HiCBrowser (https://github.com/maxplanck-ie/HiCBrowser), is also freely available to be used as a standalone browser, where users can include their own genomic tracks. With these resources, we hope to make Hi-C analysis a routine part of genomics workflows.

## Discussion

In this study, we used high resolution (DpnII restriction enzyme) and deeply sequenced (~246 million reads) Hi-C data from (Li et al. 2015; Cubeñas-Potts et al. 2016) to map the genomic positions of TAD boundaries within ~600 bp in *D. melanogaster.* We characterize TAD size, boundary strength, chromatin marks, gene orientation and transcription at the TADs. We perform motif calling at boundaries, validating the presence of known insulators, along with core promoter motif 6, motif 8 and M1BP motif, which have not been associated to boundaries before. We show that DNA motifs and open chromatin are sufficient to accurately predict a major fraction of fly boundaries. Finally, we present a set of useful tools and a resource for visualization and annotation of TADs in different organisms.

Our study verifies various properties of fly boundaries indicated in previous publications. We detect that most boundaries associate with promoters and active chromatin (Hou et al. 2012) and that various known insulator proteins are enriched at boundaries (Hou et al. 2012; Sexton et al. 2012). However, some of our results contradict previous observations. For example, we find that genes at boundaries have higher expression and low variability of expression throughout fly development in contrast with (Hou et al. 2012) who suggest that gene density and not the transcriptional state is important for boundary formation. We detect that genes at boundaries are divergently transcribed, in contrast to (Hou et al. 2012), that CTCF is not a major boundary associated insulator in Flies, in contrast to (Hou et al. 2012; Sexton et al. 2012; Van Bortle et al. 2012) and that the number of insulators at boundaries correlate very little with boundary strength, in contrast to (Van Bortle et al. 2014). Most of these contradicting results are due to two important differences of our study with previous studies:

A) *The use of higher resolution boundaries*: Most of our boundaries overlap with the known boundary protein CP190 (Fig 1B, Fig S1F) and the boundaries uniquely detected in our study have a lower TAD separation score than those unique to other studies (Fig. S1G).
B) *Analysis of combination of DNA motifs, along with ChIP-Seq peaks, rather than ChIP-Seq peaks alone*: We show that correlating boundaries with ChIP-Seq peaks alone is not a good measure when it comes to determinants of boundary formation. Many DNA binding proteins show co-localization in ChIP-Seq data, without presence of the corresponding DNA motifs (Fig. 3D, S3J). This is possible due to cross-linking artifacts and indirect binding, which is, in fact, aggravated at boundaries, which tend to contact each other in 3D space (Liang et al. 2014).

Another argument for considering Motifs is the contradicting case of CTCF at boundaries. In contrast to earlier studies (Sexton et al. 2012; Hou et al. 2012), we find that CTCF motif is rarely associated to boundaries. We could observe a significant enrichment of CTCF at boundaries in ChIP-Seq data from (Wood et al. 2011). However this enrichment disappears if we use the CTCF ChIP-seq data from (Li et al. 2015). On the other hand, if we analyse CTCF peaks containing the CTCF motif, both ChIP-seq datasets show significant enrichment (Fig. S3J). For CTCF, and in general for ChIP-seq experiments in flies, 'phantom peaks' are known to occur at active promoters (Jain et al. 2015). Thus, to avoid misleading results our analyses are based on motif presence when possible and for ChIP-Seq datasets, we try to use significance threshold along with motif binding intensity for analysis (instead of taking a significance cutoff alone).

We observe that boundary strength is affected by the chromatin states of flanking TADs and particular motif combinations, but is not affected by the number of co-occurring boundary motifs. While the boundary strength is higher between active and inactive/PcG TADs, boundaries separating two TADs within the same state (eg. active-active, inactive-inactive) are weaker (Fig. S1F). We observe that boundaries containing combinations of the motifs for Beaf-32 and either Pita, motif 6 or ZIPIC are stronger; however, the mechanism by which combinations of insulators alter the boundary strength still remains unclear. In this regard the relation between Nup98 and with both the Pita motif and motif-6 (Fig. 3D and S3H) suggest that association with nuclear pore proteins may result in stronger boundaries.

Our results indicate that the two sets of boundary motifs (promoter & non-promoter) specialize in the compartmentalization of different types of chromatin. Boundaries containing core promoter motifs are either flanking, or surrounded by active chromatin regions (Fig. 4B). In contrast, the boundaries containing non-promoter motifs tend to be within or at the borders of inactive or repressed chromatin (Fig 4B). For example, repressed TADs at Hox gene clusters are delimited by CTCF, Su(Hw) or Ibf1/2. This finding is in line with previous reports showing an enrichment of CTCF at the borders of H3K27me3 domains (Van Bortle et al. 2012; Sexton et al. 2012) and an enrichment of Beaf-32 in active chromatin (Sexton et al. 2012). An interesting speculation is that the diverse range of architectural proteins in flies have provided scope for precise control of gene regulation by allocating boundary motifs (and therefore proteins) at different places. For example, we observe that GAF motif, whose presence is negatively associated with TAD boundaries (Fig. S5 A,C), is rather detected alone at “loop domains” (Fig. S5E). It’s also important to consider the possibility that different boundary types might involve different mechanisms of formation. Furthermore, the identification of the new boundary motif 6 and motif 8 hints that there might be unidentified insulator proteins that recognize these motifs. This indicates that there is further scope to expand the list of insulator proteins in flies..

An important aspect of this study is the demonstration that TAD boundaries can be predicted with high-accuracy using motif binding affinity along with open chromatin, in absence of any other information about the presence of protein or histone marks (Fig. 5B, C). Motif binding affinity can also serve as a linear predictor of boundary strength (Fig S3B, D). These results suggest that DNA sequence is the major determinant of boundary location.

The observation that the majority of TAD boundaries are at the promoters of constitutively expressed genes favors the idea that transcription is linked to TAD formation in flies. One hypothesis, in line with the extrusion model (Sanborn et al. 2015; Fudenberg et al. 2016), is that an immobilized RNA Pol-II, anchored by the insulators, extrudes the DNA during transcription to form these domains (Fig. S6 A-B). Multiple such units working in gene-dense regions can form a rosette-like structure indicative of transcription factories (Fig. S6 C-D) (Cook 2010; Sutherland & Bickmore 2009). Promoters of divergently transcribed genes serve as good candidates for boundaries, by anchoring RNA Pol-II machines in both directions (Fig. S6 E). This mechanism, might seem more plausible in case of Flies, which have gene-dense chromosomes containing closely organized divergent promoters of housekeeping genes. However, recent observations do indicate that transcriptional activation might be linked to TAD formation in mammals (Germier et al. 2017). Further experiments would be required to visualize and quantify the relationship between transcription and TAD formation in different species.

## Methods

### Hi-C processing

Different Hi-C data available for *D. melanogaster* was downloaded from GEO and processed using the HiCExplorer (http://hicexplorer.readthedocs.io/). The following data was used:

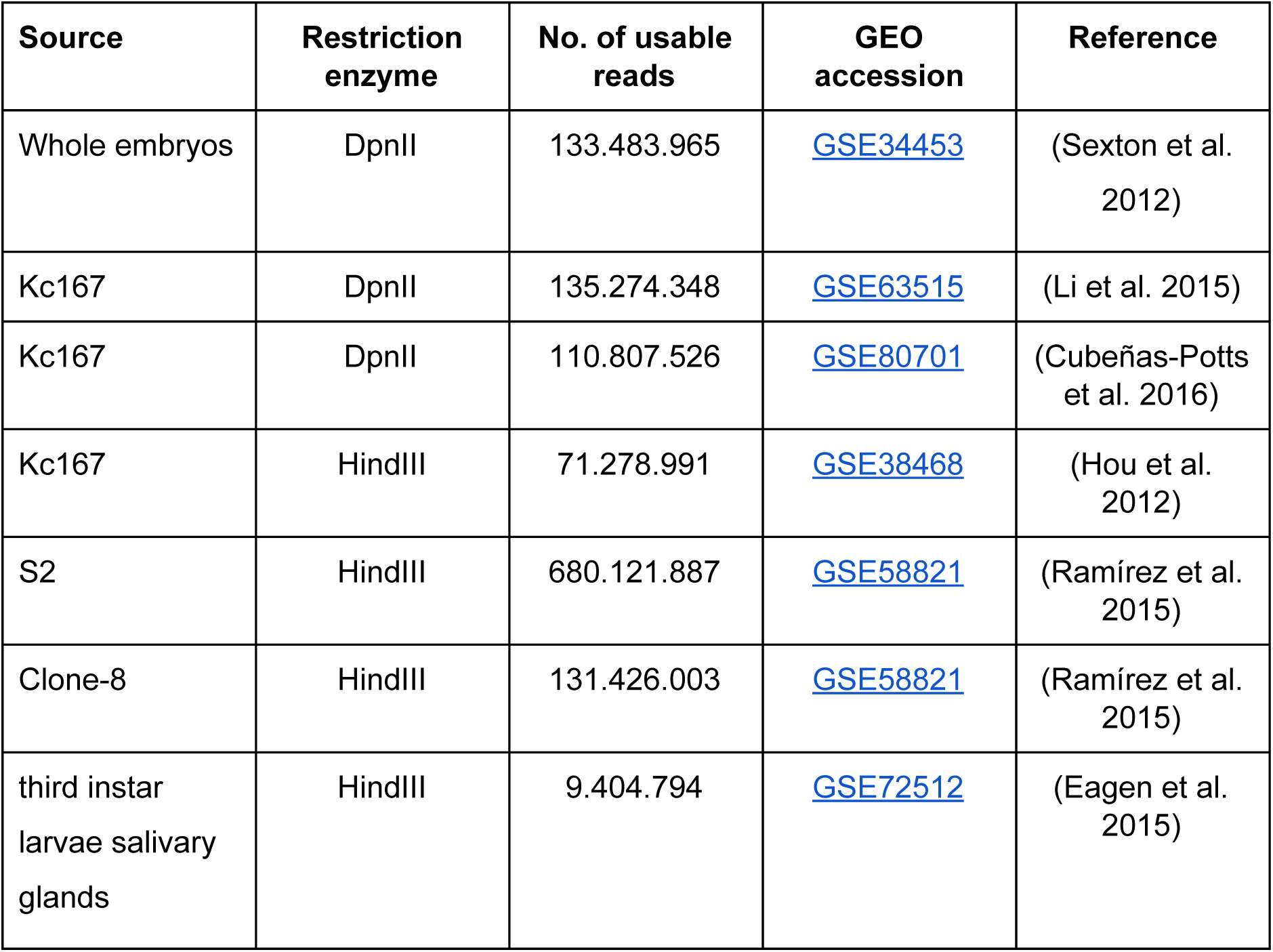

The Hi-C sequencing reads were downloaded from GEO and each mate was aligned separately using bwa mem with parameters *-E50 -L0*. The E parameter is the gap extension penalty, which is set high to avoid gapped alignments. This is because a fraction of the reads from a Hi-C experiment contain sequences from two distinct genomic positions. By increasing the gap extension penalty we promote the aligner to map the two parts of the read separately instead of trying to map the read to a single location. The L parameter is the penalty for 5’ and 3’ clipping which we set to zero to favor such clipping for the same reason as before.

To create the contact matrices, HiCExplorer divides the genome into bins of unequal size demarcated by the genomic positions of the restriction site and a matrix was created having these bins as rows and columns. The mapped reads were processed to count the number of times any two bins were connected by the Hi-C reads pairs. The following reads were discarded: read pairs that were not uniquely mapped or had a mapping score lower than 15, were within 800 bp to each other, were duplicated, contained a dangling end indicative of defective re-ligation or when one of the fragment mates was farther than 1500 bp from the restriction site.

In our processing of the data we observed that restriction enzymes do not cut with the same efficiency at all sites or sometimes do not cut at all. Because of this, after the creation of the contact matrices, rows and columns with zero or few total counts were removed. For this, we analyzed the distribution of total counts per rows. This distribution is bimodal which we interpret as two distributions combined. The distribution with lower counts contains all bins with zero reads, mostly from repetitive regions, and also the bins with low number of reads probably from inefficient digestion of restriction sites. As a cut-off to decide the minimum count of reads assigned to a bin we use the value corresponding to the valley between the two distributions. After filtering low count bins, the matrices were corrected following the iterative procedure from (Imakaev et al. 2012).

For the 4-cutter DpnII restriction enzyme the average fragment length after removing low coverage bins is 570 bp. For the 6-cutter HindIII the average was 4500 bp.

### Identification of boundaries

TAD boundaries were identified using an improved version of TAD-separation score method from (Ramírez et al. 2015) which is similar to TopDom (Shin et al. 2016). The method works by first transforming the Hi-C contact matrix into a z-score matrix **A**=(*a_ij_*). For this, each contact frequency in the matrix is transformed into a z-score based on the distribution of all contacts at the same genomic distance. For a bin *l*, the contacts between an upstream and downstream region of length *w* are in the the z-score submatrix of **A**[α_*l*_, β_*l*_], such that α*_l_*∈ {*l–w,* …, *l*} and β_*l*_ ∈ {*l*, …, *l + w*). This submatrix corresponds to the 'diamond' seen in Fig. S1A. For each matrix bin we compute the TAD-separation score (*w*) as the mean the **A** [α_*l*_, β_*l*_] values.

To reduce noise the TAD-separation score is computed for different values of *w* that are averaged afterwards per bin. Genomic bins with a low TAD-separation score with respect to neighboring regions (local minima) are indicative of TAD boundaries (Fig. S1A). To discard false positives we compare, for each local minima, the distribution of the z-score for the submatrices **Α**[α_*l*_, β_*l*_] having *l* ∈ {*i*, *i–w, i + w*}, where *i* is the bin of the local minima, and *i–w* and *i + w* are the bins at distance *w* upstream and downstream of *i* respectively. We use the Wilcoxon rank-sum test to compare the values of **A** [α_*i*_, β_*i*_] with the values of each of the other matrices and the lowest of the two *p*-values is used. Finally, we correct the *p*-values using the Bonferroni method.

To call boundaries, we used the following parameters:

*w* ∈ {10000, 12000, 18000, 25000, 40000} and *p*-value < 0.001. We also used a minimum local minima depth of 0.01. Depth of local minima (referred to as delta) can be considered similar to “fold change” of any minimum, with respect to the neighboring TAD-score average.

In contrast to other published methods to call TADs, this procedure has several advantages: i) Each boundary is associated to a TAD-separation score and a *p*-value, that are useful to characterize strong vs. weak boundaries, ii) the TAD score can be easily visualized (e. g. as an genome browser track), which is always useful for visual inspection, and iii) the computation of boundaries takes only minutes, scaling linearly with the length of the genome). Our method differs from the TopDom method in the following aspects: i) We compute TAD-separation scores using a z-score matrix while TopDom uses the corrected counts matrix, ii) we use multiple length (*w*) sizes to compute our TAD-separation score while TopDom uses a single *w* value, iii) we compute *p*-values using the 'diamond' **Α**[α_*l*_, β_*l*_] submatrices values in contrast to the 'diamond and triangle’ distributions used in TopDom. The triangle distribution contain the intra z-score values between bin *l–w* and *l*, and the intra z-score values between bin *l* and *l + w*.

### Validation of boundary quality

We used the following functional signatures to validate the quality of our boundaries :

#### Distance to known insulator co-factor CP190

Since all studied insulators proteins bind to CP190 (Ong & Corces 2014; Zolotarev et al. 2016a; Cuartero et al. 2014), a sensible quality measure is the overlap of boundaries with CP190 ChIP-seq peaks. For this, we computed the distance of the boundaries to CP190 peaks using bedtools (Quinlan & Hall 2010) *closestBed* (Fig. 1B, S1H). For comparison, we randomized our boundary positions using bedtools *shuffleBed* (Fig. S1H) and estimated the new distances to CP190. *ShuffleBed* simply assigns a new random position for each boundary anywhere in the genome (excluding heterochromatic and unplaced regions. Finally, we computed the background probability of obtaining the observed overlap between CP190 peaks and Hi-C boundaries using bedtools Fisher’s test.

#### Separation of histone marks

As boundaries are expected to separate histone marks we used the method described by (Rao et al. 2014) to quantify the correlation of marks within TADs and between TADs. For this, each TAD was scaled to 15kb, flanked with a 15kb region and divided into 1 kb bins. For each bin the mean histone ChIP-chip value was recorded, thus generating a matrix of 2,852 TADs (rows) and 45 bins (columns). For this we used computeMatrix from deepTools. The pair-wise pearson correlation value of each column was then computed to produce a matrix of size 45 × 45.

### Classification of TADs

The following histone marks for Kc167 from modEncode (Celniker et al. 2009) were used: H3K36me3, H3K4me3, H3K9me3 and H3K27me3. Other marks that correlate closely to these marks were not included. For example, H3K9me2 correlates closely with H3K9me3, H3K4me2 with H3K4me3 etc. The average intensity of the marks over the TAD length was computed using multiBigwigSummary from deepTools2 (Ramírez et al. 2016). The resulting matrix was clustered by computing euclidean distances between the histone marks and applying hierarchical clustering using the complete method. Five clusters were detected (Fig. S4B) that correspond to the presence of H3K36me3, H3K9me3, H3K4me3, H3K27me3 or none. Analysis of the TADs containing H3K36me3 and H3K4me1 in the genome revealed that that H3K36me3 is present at exons of active genes while H3K4me1 is mostly present at introns and intergenic regions of active genes and less abundant at exons. Thus, noticing that these two marks are complementary for active regions we classified TADs having predominantly these marks as ‘active’. For the other clusters we used the same categories as (Filion et al. 2010): the cluster of TADs with H3K9me3 was labeled as ‘HP1’ (Heterochromatin Protein 1); the cluster with H3K27me3 was labelled ‘repressed’ or ‘PcG’ (Polycomb group) and the cluster with no mark was labelled as inactive.

### Analysis of transcription at boundaries

In order to analyse transcription at boundaries, we downloaded ribo-depleted RNA-Seq data from modENCODE (Celniker et al. 2009). Data was mapped to the Drosophila (dm3) genome using HISAT2 (v2.0.4) (Pertea et al. 2016) and the reads were summarized per gene using featureCounts (v1.5.0.p1) (Liao et al. 2014) using options *-p --primary -Q 10*. We used data from Kc167 cells, along with 14 different developmental stages, ranging from embryo to adult. We only used data produced in 2014 in order to avoid batch effects and further confirmed the data quality by clustering the samples by euclidean distance (Fig S2A). We normalized each sample by library size, averaged the counts for replicates, and finally used log transformed counts for all the analysis.

Genes were considered expressed if they have a normalized log-count of 1 or more. Variability was assessed using coefficient of variation of a gene across all developmental stages along with Kc167 cells. For gene-pairs in any orientation (convergent, divergent or tandem) we calculated the correlation (spearman and pearson) of coef. of variation of gene-1 with gene-2 across development and plotted the results. We further tested whether overall, genes within TADs tend to be more correlated in expression than genes between TADs. For this we used a subset of consecutively arranged TADs that have more than one gene inside them. We then used ANOVA to test for differences between and within TADs, as seen in Fig S2C, many TADs pairs are significantly different from each other, while very few TADs are significantly different if we randomly assign genes to TADs.

### Identification of boundary motifs

We took the list of boundaries and expanded them by 500 bp on each side. To avoid false positives, repetitive regions from the sequences of those boundaries were replaced by ‘N’s and any region with more than 10% of ‘N’s was removed. We used MEME-chip (Ma et al. 2014) to identify enriched DNA motifs; MEME-chip internally computes motifs using two methods, DREME (Bailey 2011) and MEME (Bailey & Elkan 1994). DREME aims to quickly identify short motifs while MEME identifies larger overrepresented sequences (at the expense of significantly longer processing times). We used the consensus of DREME and MEME to call motifs.

To obtain the position-weight matrices of insulator motifs we ran MEME-chip (Ma et al. 2014) on the peaks called using MACS2 (Feng et al. 2012). We selected the highest scoring motif for each case which invariably corresponded to the motif reported for the protein. We used ChIP-Seq data for Beaf-32, CTCF and Su(Hw) from (Wood et al. 2011) GAF from (Cubeñas-Potts et al. 2016); Ibf1/2 from (Cuartero et al. 2014); Pita and ZIPIC from (Maksimenko et al. 2015); and Zw5 from (Gaszner et al. 1999).

### Enrichment of motifs using control background

For promoter boundaries, a control background composed of all drosophila gene promoters was used to test the enrichment. We downloaded drosophila genes (dm3 assembly) from UCSC table browser (Karolchik et al. 2004) and selected the sequences 200 bp upstream and 50 bp downstream of the transcription start site as core promoter sequences.

We classified these promoter sequences as boundary if they were within 500 bp of a boundary, or non-boundary (control background) if they were farther than 2,000 bp from a boundary. Repetitive regions from the sequences of those promoters were replaced by ‘N’s and any region with more than 10% of ‘N’s was removed. In total, 10,529 background promoters and 1,944 boundary promoters were used.

We used two different methods to assess the enrichment of the *de-novo* and known motifs in boundary promoters with respect the control background, namely Ame (McLeay & Bailey 2010) from the MEME-suite and a method based on the predicted binding affinity given by TRAP (Thomas-Chollier et al. 2011) that works as follows: for each motif, the log(TRAP score) distribution was computed for both the background and the boundary promoters. The Wilcoxon rank-sum test was then used to test for differences in the distributions. The P-values obtained were corrected using FDR. For Ame we use total hits as scoring method and Fisher’s test for estimating enrichments. We tested all *de-novo* motifs identified either by MEME or by DREME and all known motifs associated to insulators and CP190 cofactors as well as all core-promoter motifs.

We also used as control active genes in Kc167. To make this control, we selected those genes that overlapped with the yellow and red chromatin states from (Filion et al. 2010) that are indicative of active chromatin in Drosophila Kc cells. The enrichment results were similar to the ones using a more broader list of genes for background.

For the boundaries that are not at promoters we used non-promoter open chromatin sequences obtained from DNase-seq from (Celniker et al. 2009) as control. In this case we used 1,665 background open chromatin regions and 655 non-promoter boundaries.

### Pausing index

Pausing index for all *D*. *melanogaster* promoters was computed as the ratio of Pol-II ChIP-seq coverage at promoter over coverage at gene body. We used the ChIP-seq data for RNA Pol-II from (Li et al. 2015). The promoter region was defined as in the previous section (200 bp downstream, 50 bp upstream of transcription start site). The gene body was defined as the region between 50 bp downstream of the transcription start site and the gene end. We used the maximum coverage for the promoters and the median coverage for the gene body.

### ChIP-seq data sources

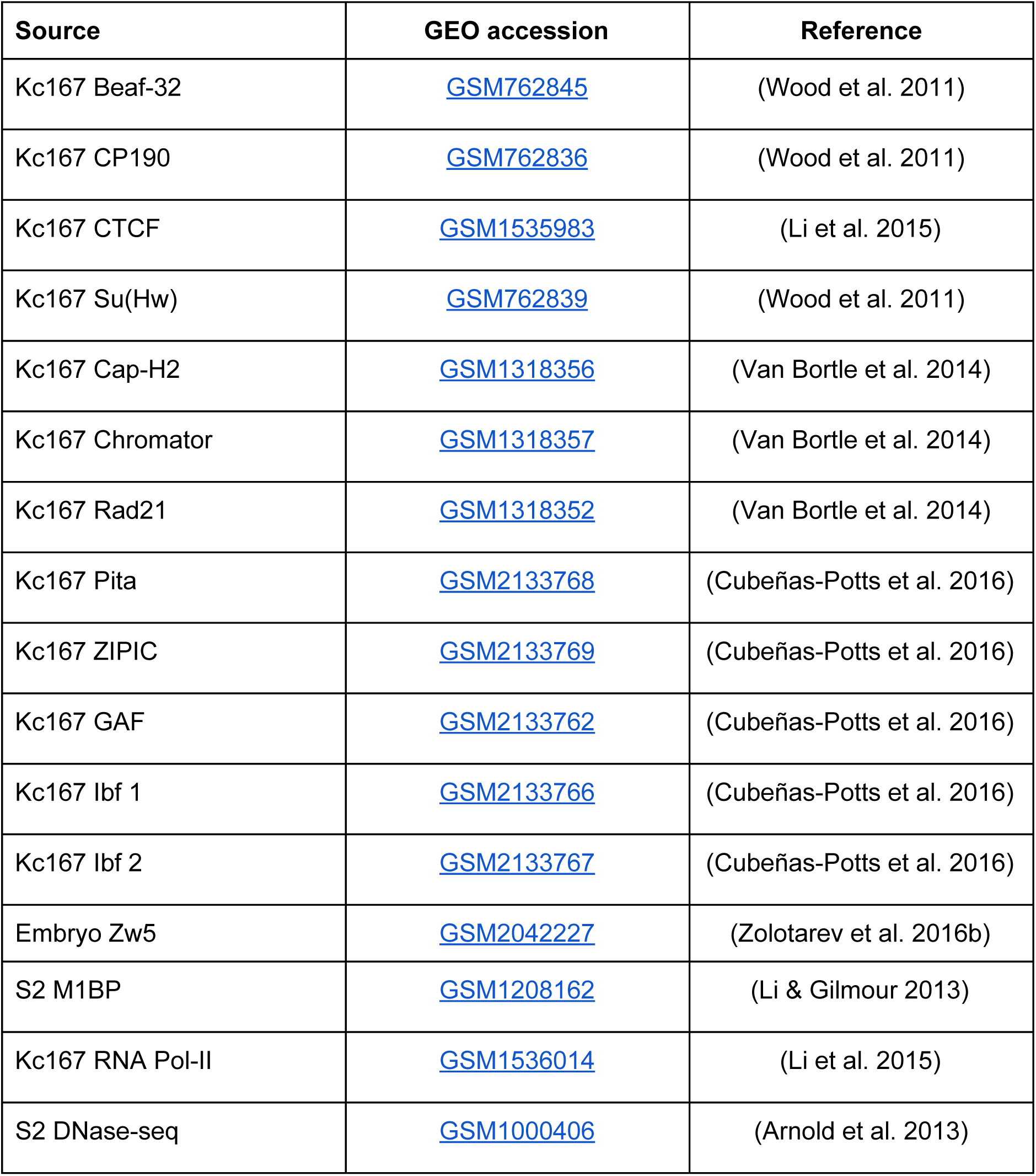

### Processing of ChIP-seq data

For each ChIP-seq data used, we downloaded the respective fastq files and aligned them in the dm3 assembly using Bowtie2 (Langmead & Salzberg 2012). MACS2 was used to identify peaks for each of the proteins (Feng et al. 2012). For the respective data sources we downloaded input sequences and aligned them as the ChIP-seq data. *bamCompare* and *bamCoverage* from deepTools2 (Ramírez et al. 2016) were used to create normalized coverage bigwig files.

MEME-chip (Ma et al. 2014) was used to identify motifs based on the MACS2 peaks. The resulting motifs can be seen in Table 1.

### Clustering of motifs

We used the promoters (200 bp upstream 50 bp downstream) annotated as boundaries and computed the log(TRAP score) for the Beaf-32 motif, motif-1 (M1BP), motif-6, motif-7 (ZIPIC) and motif-8. The scores for each motif were converted to bigwig files and clustered using hierarchical clustering from deepTools2 (Ramírez et al. 2016).

All boundaries that were further than 2000 bp of a promoter were centered at the nearest CP190 ChIP-seq peaks within 2000 bp, otherwise the boundary position was not modified. Log(TRAP score) for CTCF, Ibf and Su(Hw) were computed for these regions and clustered as previously described.

We used hierarchical clustering based on euclidian distance and the ward method. The cluster number used was 13 for promoter boundaries and 9 for non-promoter boundaries. In each case, the group compose only low TRAP scores was removed. After clustering, the groups were manually ordered and to produce the left panel of Fig. 2C. Scale of each heatmap was manually adjusted based on the range of TRAP scores found at the clusters for each motif (Fig. S3A). The log2 ratio of ChIP-seq / input for the different proteins was used for the center and right panels of Fig. 2C. Each heatmap is centered on the boundary and extended +-5000 bp. Scale of the heatmaps were adjusted based on the log2 ChIP/input for the protein in the respective cluster (Fig. S3B).

### Motif presence

For figure 3, we considered a motif as present at a boundary if the TRAP score was equal or higher than the minimum log(TRAP score) identified for the clusters in Fig. 3C (the distribution of the log(TRAP scores) can be seen in Fig. S3A). The thresholds used were: ZIPIC motif -4.7, Beaf-32 -5, M1BP -4.5, motif-6 -3, motif-8 -2, Ibf -4, CTCF -4, Su(Hw) -3. For GAF, Pita and Zw5 motifs we used FIMO (Grant et al. 2011) with the following parameters: *‘--max-strand --thresh 1e-3’.* For analysis of motif combinations at boundaries, we also require that the motifs are accompanied by the corresponding ChIP-seq peaks. For motif-6 and motif-8 whose binding proteins are not known, we require that the motif is on an accessible region. For this we use the peaks from the DNAse-seq data (Celniker et al. 2009).

### Boundary prediction and feature ranking

We performed boundary prediction at all drosophila promoters using motif TRAP scores for various transcription factors and DNAse-Seq signals as features. We utilized methods ranging from simple to complex (linear models, logistic regression, random forest and stochastic gradient boosting), with the primary purpose to rank the features by importance in boundary prediction. Pre-filtering was done to remove highly correlated features (pearson R > 40%). Linear model and random forest was performed using the package *Caret* (<u>Kuhn et al. 2016</u>), while logistic regression was performed using package *glmnet* (Friedman et al. 2010) in R.

Linear model was used with stepwise feature selection algorithm to predict boundary score from features by selecting the combination of features that minimizes the Akaike Information Criteria (AIC). Logistic regression, Random forest and gradient boosting were used to classify the promoters into boundary and non-boundary, with additional feature selection performed using lasso, for logistic regression. We performed 10-fold cross validation while training all classification models. To evaluate model accuracy, the data was randomly divided into training (60%) and test (40%) datasets and the sensitivity and specificity was calculated for test predictions. Lasso and gradient boosting models show highly similar sensitivity and specificity when used on new test dataset, compared to when same dataset was used for prediction, suggesting they are robust and less prone to overfitting.

Linear model predicted the boundary scores on the test dataset with overall spearman correlation of 37.6%, while logistic regression and random forest performed predictions with around 73 to 78% accuracy. After obtaining the best model in each scenario, we ranked the features by their importance in prediction, using the *varImp* function from *caret*. *varImp* selects a variable importance predictor based on the model type, which is calculated for each parameter in the model (https://topepo.github.io/caret/variable-importance.html). Briefly, the importance score for linear model is the absolute value of the *t*-statistic for the model parameter, for lasso, it’s the absolute value of final coefficients, for gbm it’s the relative influence score as described in (Friedman 2001), and for random forest it’s the difference between the classification error-rate for the out-of-bag portion of data and a permuted predictor variable, averaged over all trees and normalized by the standard deviation of the differences (https://www.stat.berkeley.edu/~breiman/RandomForests/cc_home.htm#varimp). All importance scores are then scaled between 0 and 100 to compare them together.

## Supplementary Material

- Supplementary Table S1: List of promoter boundaries with annotations for TAD-separation score, motif and ChIP-seq enrichment.
- Supplementary Table S2: List of non-promoter boundaries with annotations for TAD-separation score, motif and ChIP-seq enrichment.
- Supplementary Table S3: List of TADs annotated with their classification.

## Acknowledgments

We would like to thank Wilhelm Rüssing for IT support and Diana Santacruz for critical reading of the manuscript. We also thank Victor Corces lab whose published data has been instrumental in this research.

## Funding

This work was supported by the German Research Foundation [SFB 992, Project Z01] and a grant from the Federal Ministry of Education and Research through the German Epigenome Programme DEEP [01KU1216G]. Source of Open Access funding: own funds. Conflict of interest statement: None declared.

